# Need for cognition moderates cognitive effort aversion

**DOI:** 10.1101/2023.06.20.545770

**Authors:** Davide Gheza, Wouter Kool, Gilles Pourtois

## Abstract

When making decisions, humans aim to maximize rewards while minimizing costs. The exertion of mental or physical effort has been proposed to be one those costs, translating into avoidance of behaviors carrying effort demands. This motivational framework also predicts that people should experience positive affect when anticipating demand that is subsequently avoided (i.e., a “relief effect”), but evidence for this prediction is scarce. Here, we follow up on a previous study (1) that provided some initial evidence that people more positively evaluated outcomes if it meant they could avoid performing an additional demanding task. However, the results from this study did not provide evidence that this effect was driven by effort avoidance. Here, we report two experiments that are able to do this. Participants performed a gambling task, and if they did not receive reward they would have to perform an orthogonal effort task. Prior to the gamble, a cue indicated whether this effort task would be easy or hard. We probed hedonic responses to the reward-related feedback, as well as after the subsequent effort task feedback. Participants reported lower hedonic responses for no-reward outcomes when high vs. low effort was anticipated (and later exerted). They also reported higher hedonic responses for reward outcomes when high vs. low effort was anticipated (and avoided). Importantly, this relief effect was smaller in participants with high need for cognition. These results suggest that avoidance of high effort tasks is rewarding, but that the size off this effect depends on the individual disposition to engage with and expend cognitive effort. They also raise the important question of whether this disposition alters the cost of effort per se, or rather offset this cost during cost-benefit analyses.

## Introduction

Daily life constantly presents us with challenging tasks that we need to perform to achieve our goals. Successful completion of these tasks requires us to invest cognitive effort. Recent theories (2,3) suggest that such decisions are implemented as simple cost-benefit analyses, where reward is discounted by the anticipated cost of effort. The idea that effort is costly is captured by the ‘law of least effort’ (4). This law predicts that, all else being equal, people will tend to avoid effortful actions. There is ample empirical evidence for this principle. For example, Kool and colleagues (5) demonstrated that human participants prefer choice options with the least cognitive demands (even when time on task and error likelihood were matched). Such effort avoidance has been demonstrated for a wide range of demands, such as response conflict (6,7), task switching (5,8), complex task policies (9), and short-term memory load (5). In fact, a recent study demonstrated that people are willing to endure physical pain to avoid effort (10). Across these studies, the task demands invoke the need for cognitive control or controlled information processing (11,12), which are core features in models of mental effort (13). In short, mental effort carries a cost, and this cost is used to devaluate anticipated reward (14–16), resulting in an expected value of control’ (2).

If reward is discounted by effort, then it should also be enhanced if effort is anticipated, but eventually avoided. That is, the avoidance of expected effort should be experienced as relief. Interestingly, relief (from anticipated pain) appears to involve the same reward processing systems that underly representations of effort costs (17,18). Together with the observation that effort is costly, this predicts that the avoidance of anticipated effort should be experienced as rewarding. To our best knowledge, there is a scarcity of studies that investigate this phenomenon. Here, we report two studies that test this hypothesis.

Earlier work from our group (1) has provided some initial, but ultimately inconclusive, support for this hypothesis. In this experiment, participants performed a well-validated gambling paradigm (19) that was combined with an effort task (20). On each trial of this task, participants chose between four options that each afforded a chance to win a reward. Critically, on some of the trials, participants were informed that after a loss (a ‘no reward’ outcome), they would be given the choice to exert effort to repeat the gamble. Therefore, on these ‘special’ trials, there was a prospect of effort both prior to and during gambling, while for the rest of trials, there was not. Moreover, rewards on special trials were associated with the avoidance of effort (i.e., it rendered the opportunity to spend effort to repeat the gamble moot). Interestingly, the participants reported more positive affect (pleasantness), as well as increased relief for reward delivered on special trials compared to standard trials. In addition, at the electrophysiological level, we found that reward delivered after gambling on special trials elicited a larger reward positivity event-related potential (21) as well as enhanced power in delta (22) and beta-gamma (23–25) frequency bands. Together, these results suggest that relief from effort registers as rewarding (4,26,27). However, an alternative interpretation exists.

To see this, note that people did not just avoid mental effort on rewarded special trials, but they also avoided the extra time spent on deciding whether to repeat the gamble and on the effort task itself. This creates an opportunity cost (28): special trials with no reward lasted longer, and thus were associated with reduced time to accrue reward. In other words, participants may have experienced more positive affect for rewards on special trials simply because it reduced their time on the experiment. Thus, though these results are suggestive, it remains to be shown whether relief from effort enhances subjective reward.

To test this, we made several key changes to our experimental design (1), so that only demands for mental effort, and not opportunity costs, differed between conditions. In this new task, each trial with no reward was followed by a cognitive effort task (29) performed to recuperate the missed reward. This cognitive effort task required mental arithmetic. At the start of each trial, a cue indicated the amount of demanded effort for the arithmetic task (i.e., easy vs. hard). Comparing self-report ratings between these trials allowed us to isolate the effect of effort avoidance, whereas in our previous study the analogous comparison conflated effort and time. It is worth drawing attention to the fact that each trial with no reward lead to the effort task. Participants did not decide whether or not to perform the task. Therefore, the expectation of effort was high and constant across all trials. These methodological changes allowed us to compare the subjective experience of reward when either an easy or difficult task was expected, with opportunity cost equal between conditions.

The second goal of our study was to assess the source of individual differences in effort-related relief. In addition to objective task demands, valuations of effort are influenced by people’s state and trait dispositions. For example, stress increases effort avoidance (30) and depression/anhedonia decreases reward processing (31). Here, we tested whether relief was predicted by participants’ ‘need for cognition’ (NFC; (32)). The NFC is defined as the tendency to engage in and enjoy effortful cognitive activities (e.g., those requiring thinking and problem solving; see also (33)) and can be conceived as a motivational drive (34). We hypothesized that an increased NFC would decrease the relief of effort avoidance, and hence behave in an opposite manner compared to stress, depression or negative affect (35). In other words, the NFC might mitigate the impact of effort (anticipation) on reward processing, with a relatively reduced effort-related relief for participants with a high NFC.

## Methods

### Participants

Twenty-three undergrad students from Ghent University (17 females; median age: 21 years, range: 18-30) participated in Experiment 1. They had normal or corrected-to-normal vision and did not report any history of neurological or psychiatric disorders. Sample size was determined to be at least as large as in our previous experiment (1) where a similar experimental manipulation was used, and where a significant effect of cost anticipation on reward was found (increased pleasantness [p = .008, d = 0.57] and relief [p = .033, d = 0.27] of reward when cost could be avoided (1).

Seventy-nine young adults participated in Experiment 2. None of them participated in Experiment 1. To explore individual differences in NFC, the sample size was determined to be as large as available time and resources would allow. Three participants were excluded from further analyses due to low precision in their ratings of the reward-related feedback (see exclusion criteria below). The final sample consisted of 76 participants (62 females; median age: 21 years, range: 18-46).

These two experiments were part of a more general research project investigating effects of motivation on reward that was approved by the local ethics committee at Ghent University. All participants gave written informed consent prior to the start of the experiment, were debriefed at the end, and received a monetary compensation for their participation. Participants’ data was collected between 2018 and 2019 and was anonymized at the time of collection.

### Stimuli and task

For Experiment 1, we adapted a widely used gambling task (1,36) and combined it with a cognitive effort task (29). At the start of each trial, participants were informed about the cognitive effort level with a text cue (“easy” vs “hard”) located at the center of the screen (1000 ms). Following a fixation dot (1500 ms), four doors appeared on the screen, and participants had to choose one of them by pressing with their left hand the corresponding numeric key (1 to 4) on a keyboard. After another fixation dot (700 ms), this choice was followed by reward-related feedback (1000 ms), indicating either a reward (green “+”) of 6 cents, or a no-reward outcome (red “o”). Participants were instructed to guess and select a door containing a reward in order to maximize their payoff. However, unbeknown to them, the outcome was unrelated to the choice and reward probability was set to exactly 50%. Participants were instructed that in case of a reward, the trial would end and a new trial would follow. Hence, no additional effort would be necessary. However, when there was no reward, a second task would follow, which could be hard or easy (as indicated by the previous cue), so that they could get another chance to win reward. Thus, receiving no reward during the gambling task resulted in the prospect of effort. More specifically, after 1000 ms (fixation), a mental arithmetic task started (i.e., the ‘effort task’). This task required participants to complete two calculations (two additions or an addition and a subtraction, all of which with single-digit numbers) (see Figure 1). In the hard condition, every operation required carrying or borrowing. In the easy condition, none of the two operations required carrying or borrowing. This manipulation results in two levels of difficulty, as shown in previous studies (37–39) and confirmed by subjective ratings (see results below).

**Fig 1.**
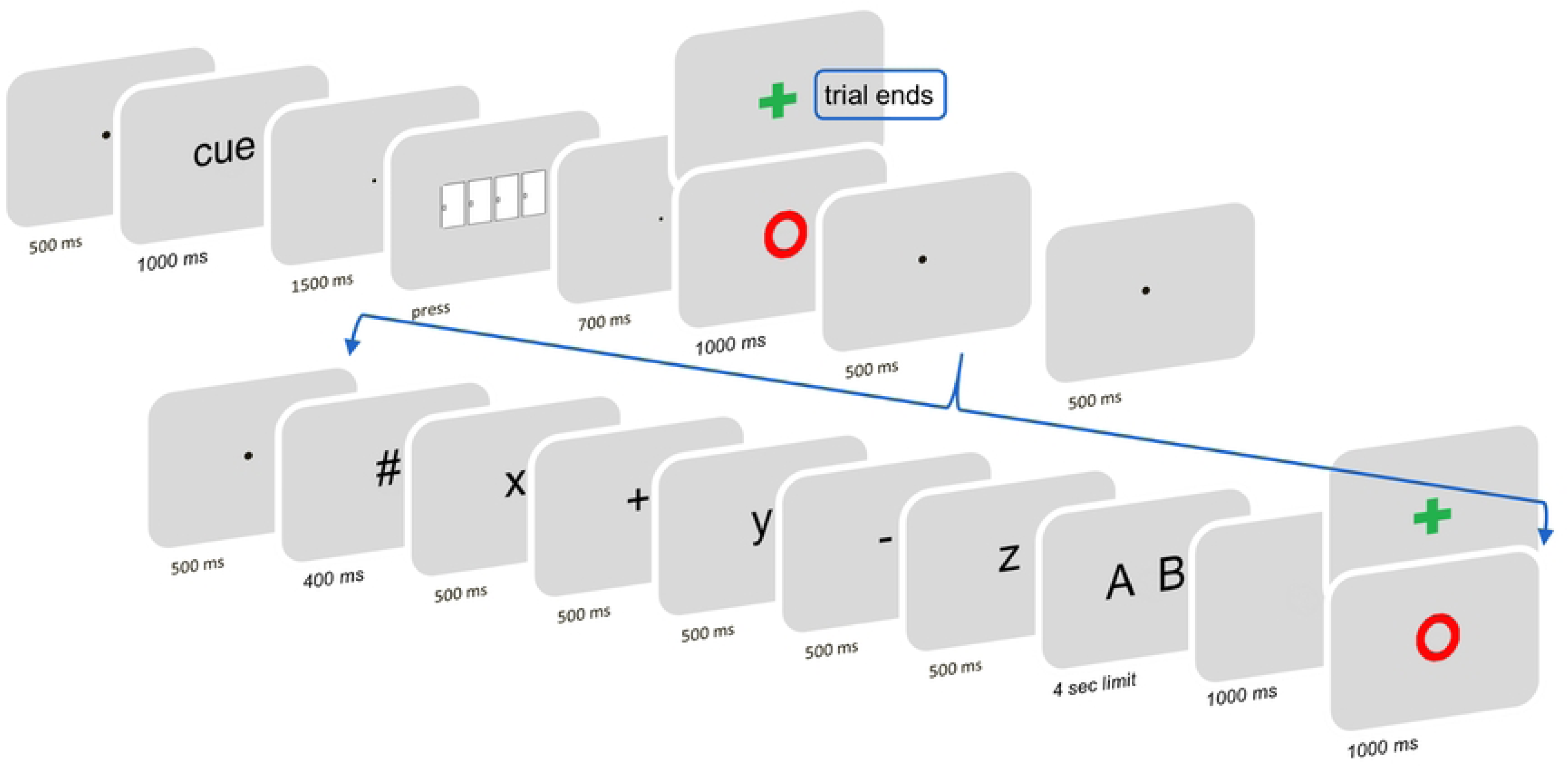
Overview of the task and trial structure. Participants were first informed about the cognitive effort level with a text cue (word “easy” or “hard”). After they picked one door, they received reward or no-reward feedback (50% reward probability). Only in case of no-reward feedback, the effort task ensued. For both conditions, the initiation of the effort task was probed by an uninformative pound symbol (#), followed by a serially-presented list of digits and operations. Participants were instructed to integrate them, and based on this arithmetic select the correct one out of two possible answers. Based on their answer, “performance” feedback was given, providing reward in case of a correct answer. On a small proportion of trials, the subjective hedonic value of the reward-related feedback was assessed with probes presented 1000 ms after its offset.

The effort task was structured as follows: a pound symbol indicated the start (400ms); digits and arithmetic signs were then presented serially, each lasting 500ms and interleaved with blank screens (200ms in Experiment 1, 100ms in Experiment 2); finally, two possible solutions were presented simultaneously, and the participants had to choose the correct one by pressing the corresponding key with the right hand (i.e. numeric keypad; “1” for the leftmost or “2” for the rightmost solution). They were instructed to select the correct answer as quickly as possible, with a time limit of 4000ms. After this choice, a blank slide was presented (1000 ms), followed by a new feedback screen related to their performance (1000 ms): a reward outcome (green “+”) indicated a correct response and a win of 6 cents, and a no-reward outcome for incorrect responses (red “o”) was presented. If participants missed their response, or if they responded too late, a screen indicated that there was “no response detected”. Serial presentation of digits and operations was chosen in order to equate the time spent observing stimuli between conditions, as well as to avoid calculation strategies (29). Across trials, different combinations of digits and arithmetic signs were used to avoid learning or habituation. After the effort task, a new trial of the gambling task followed. The intertrial interval was fixed and set to 1000 ms.

The subjective value of the first feedback screen (after gambling) was assessed by specific probes. In a few trials (n = 48), 1000 ms after the offset of the gambling outcome, three questions were presented, probing the perceived pleasantness, frustration, and relief of the outcome. Participants answered them using visual analog scales (VAS). These three ratings were submitted 12 times for each effort level (low vs. high) and outcome (reward vs. no reward) condition.

Experiment 1 consisted of 208 trials, including an equal amount of easy and hard trials. The gambling task had a pre-set reward probability of 50%, and the effort task had to be completed only in case the choice did not deliver reward. Therefore, the effort task was administered 52 times for each difficulty level.

Seven self-paced breaks were distributed equally throughout the experiment. At the end of each break, participants were asked to rate the difficulty of the effort task, its pleasantness, their motivation to complete it, and their satisfaction with correct performance on the task. Each question was submitted twice, for each of the two difficulty levels, using a VAS. Participants received a fixed €8 compensation for their participation. Depending on their accuracy with the effort task, a maximum payoff of €12,48 could be earned (mean = €11.91).

For Experiment 2, where the NFC questionnaire was also administered (40), the same procedure was used but a few changes were made. The reward feedback with the gambling task indicated that 5 cents were won (instead of 6). The three ratings about the gambling outcome were submitted each 40 times in total (instead of 48), 1000ms after its offset (10 times for each combination of difficulty level and outcome). In Experiment 2 we also probed the subjective value of the performance feedback (after the effort task). Analogous to the reward-related feedback, three ratings probing the perceived pleasantness, frustration, and relief of this feedback. These were presented directly after the performance feedback. Performance feedback ratings were submitted up to 26 times, equally split between high and low effort, but were omitted in case of an incorrect response. At this stage we only probed affective ratings for correct feedback as the effort task was tuned to induce only very few errors. Reward-related feedback (gambling task) and performance feedback (effort task) were never probed in the same trial. Experiment 2 consisted of 200 trials, with an equal amount of easy and hard trials (100 each). For each of these, there were 50 trials where the effort task had to be performed (no reward) and 50 trials where it was avoided (reward). Participants were compensated with €8 for their participation. Depending on their accuracy with the effort task, they could earn up to €10 (mean = €9.63). Last, in Experiment 2 we added 16 “catch trials” to promote and assess participant’s attention to the effort cues presented at the beginning of the trial. More precisely, after door selection participants were asked to report how hard a following effort task would be. After their response, participants received catch-related feedback (correct, incorrect, or too late response) and the catch trial terminated. Eight catch trials, for each of the two effort levels, were randomly interspersed among the 200 task trials.

The experiments’ duration was approximately 60 minutes, including instructions and a short practice. In Experiment 2, the NFC questionnaire was administered after the task. All stimuli were shown against a grey homogenous background on a 21 inch CRT screen and controlled using E-Prime (41).

### Data analysis

We evaluated the effectiveness of the cognitive effort manipulation by comparing performance on the effort task (i.e., accuracy and speed) between the easy and hard conditions. Moreover, we also compared their subjective value by analyzing the ratings of difficulty, pleasantness, motivation to perform well, and pleasure in performing well. These ratings were first transformed to percentages, setting anchors to the boundaries of the scales, and were averaged across the seven repetitions.

The subjective ratings of the reward-related feedback (the VAS scores) obtained for each difficulty level (easy vs. hard), outcome (reward vs. no reward), and affective dimension (pleasantness, frustration, and relief) were also first transformed to percentages, setting anchors to the boundaries of these scales. For the “frustration” scale, we reverse-scored the percentages in order to provide comparable ratings for the three affective dimensions (all going from negative to positive).

For Experiment 2, subjective ratings of the performance feedback were analyzed in the same way. Additionally, for statistical analyses the VAS scores and NFC data were centered by subtracting the group means from the individual scores.

Participant’s data were excluded from further analysis if the average mouse-click x-coordinate of the reward-related feedback rating was above 105% or below -5% of the rating range, in any of the 3 affective dimensions, and any of the 4 levels of outcome by difficulty.

### Statistical analyses

Subjective ratings data were analyzed using Bayesian model comparison. Inference about their generative processes was based upon Bayes Factors (BFs), computed for alternative explanatory models in ANOVA designs (41, see also 42). The analyses’ pipeline was implemented in R v4.0.5 (44) with the package BayesFactor v0.9.12-4.2 (45), and involved: I) defining theoretically sound probability models; II) computing BFs, i.e. the ratio between the likelihood of each model of interest (the probability of the observed data, given the model/hypothesis) and the likelihood of the null model; III) model selection based on the highest BF; IV) characterizing the direction of follow-up contrasts by means of Bayesian t-test between conditions of interest. The models’ likelihood was estimated using Markov-Chain Monte Carlo simulations with 10,000 iterations, and BFs were computed assuming a wide Cauchy prior centered on zero: d ∼ Cauchy (0, 0.707).

For the gambling task feedback ratings, the models of interest included the effects of 1) *outcome*, 2) *difficulty level*, 3) *outcome + difficulty level*, and 4) *outcome x difficulty level*. Additionally, for Experiment 2 we tested a model including 5) the three-way interaction of outcome x difficulty level x NFC score. The null model was a simple intercept model. In addition, the effects of random factors *participants*, *affective dimension*, and their interaction were specified as nuisance in all the models, including the null model.

To analyze performance feedback ratings (Experiment 2), we used a one-tailed Bayesian t-test to estimate the evidence in favor of increased positive evaluation of a reward outcome after high vs. low effort, as compared to a null model.

## Results

### Experiment 1

Accuracy on the effort task was higher for the easy (M = 98 %, SD = 14) compared to the hard condition (M = 87 %, SD = 34; BF+0 = 2.20 x 10^3^). Mean reaction time was larger for the hard (M = 1091 ms, SD = 476) compared to the easy condition (M = 621 ms, SD = 144; BF+0 = 4.11 x 10^3^). These results indicated that the difficulty manipulation was successful.

The subjective ratings of the effort task indicated these two difficulty levels were experienced as clearly different. The hard compared to easy condition was perceived as more difficult (M easy = 6.1, SD = 7.3; M hard = 31.2, SD = 18.7; BF-0 = 1.09 x 10^5^) and less pleasant (M easy = 83.4, SD = 12.5; M hard = 62.7, SD = 21.0; BF+0 = 1.92 x 10^3^), while participants reported similar levels of motivation to perform them correctly (M easy = 85.1, SD = 14.8; M hard = 86.6, SD = 13.6; BF01 = 3.58), as well as pleasure in performing them correctly (M easy = 81.8, SD = 17.1; M hard = 76.4, SD = 18.5; BF01 = 2.00).

These reward-related feedback ratings (Figure 2) were best explained by a model that contained the *outcome* x *difficulty level* interaction. Under this model the observed data were BF10 = 5.62 x 10^648^ times more likely to be produced than under the null model. Moreover, the *outcome* x *difficulty* model explained the observed data 325 times better than the second-best model, which only included the main effect of *outcome*. Follow-up Bayesian one-tailed t-tests showed strong evidence for the hypothesis that no-reward outcomes were evaluated as more positive in low-effort compared to high-effort trials (BF+0 = 6.00 x 10^5^). Conversely, for reward outcomes, there was only weak support (BF+0 = 2.22) for more positive evaluations in the high-compared to low-effort trials. In other words, participants rated a no reward outcome as more positive when they anticipated low effort. For either a reward (top) or no-reward (bottom) outcome, light grey density shades correspond to low effort anticipation and dark grey density shades correspond to high effort anticipation. The horizontal line represents the median, the box represents the interquartile range, and the whiskers extend to the last data point within 1.5 times the interquartile range.

**Fig 2.**
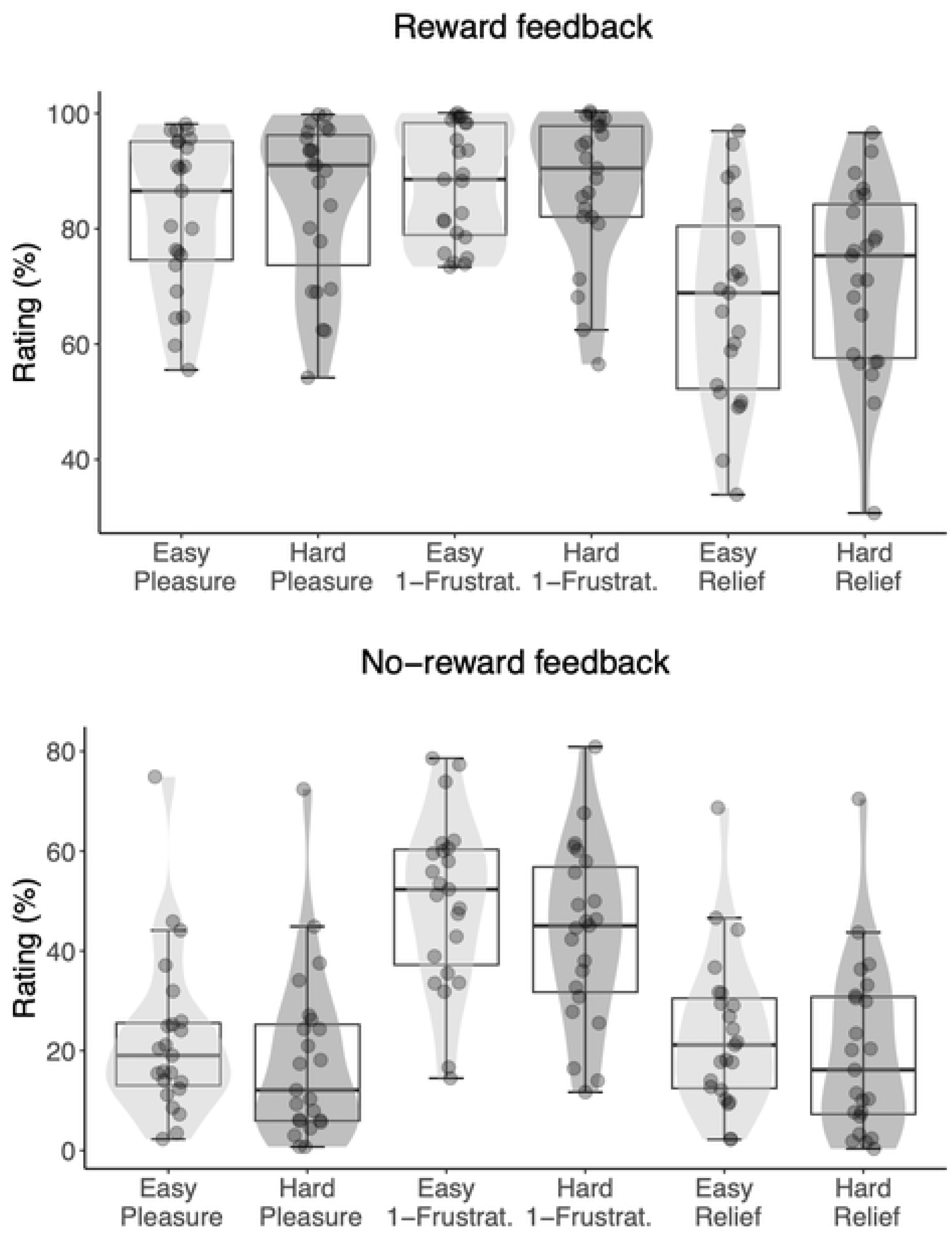
Experiment 1. Ratings of reward-related feedback that followed the gambling task, by difficulty level (easy or hard), and affective dimension (pleasure, reverse-scored frustration, and relief). Each dot represents the subject-level average across 12 repetitions of the rating.

### Experiment 2

Participants paid attention to the effort cues presented at the beginning of each trial, as suggested by high accuracy in reporting them during catch trials (M easy = 96.2, SD = 8.9; M hard = 91.8, SD = 13.8).

Similarly to Experiment 1, the accuracy for the effort task was higher for the easy (M = 99 %, SD = 12) compared to the hard condition (M = 86 %, SD = 34; BF+0 = 3.23 x 10^16^). Mean reaction time was larger for the hard (M = 1267 ms, SD = 467) compared to the easy condition (M = 694 ms, SD = 198; BF+0 = 7.21 x 10^19^). This demonstrated that the difficulty manipulation was successful.

The subjective ratings of the effort task again indicated these two difficulty levels were experienced as clearly different. The hard compared to easy condition was perceived as more difficult (M easy = 14.0, SD = 15.7; M hard = 40.2, SD = 23.1; BF-0 = 4.39 x 10^14^) and less pleasant (M easy = 76.7, SD = 19.9; M hard = 57.0, SD = 21.7; BF+0 = 6.46 x 10^7^). In line with Experiment 1, participants reported similar levels of motivation to perform the easy and hard tasks correctly (M easy = 82.0, SD = 16.1; M hard = 84.5, SD = 15.7; BF01 = 2.01), and also similar pleasure in performing them correctly (M easy = 76.3, SD = 19.3; M hard = 79.1, SD = 17.8; BF01 = 3.39).

Next, we turned our attention to the analyses of subjective ratings as a function of both gambling outcome and difficulty (Figure 3). Those ratings were again best explained by a model that included the *outcome* x *difficulty level* interaction. Under this model, the observed data were BF10 = 9.06 x 10^959^ times more likely to be produced than under the *null* model. This model also explained the observed data 1.63 x 10^14^ times better than the second-best model, which only included the main effect of *outcome*. The follow-up Bayesian one-tailed t-tests showed strong evidence for the hypothesis that no-reward outcomes were evaluated as more positive on low-effort compared to high-effort trials (BF+0 = 1.55 x 10^14^). Conversely, for reward outcomes, there was strong evidence that evaluations were more positive in the high compared to low effort trials (BF+0 = 2.08 x 10^5^). In sum, we replicated Experiment 1: participants rated the no reward outcomes as more positive when they anticipated low effort. In extension of Experiment 1 (Figure 2), participants also rated the reward outcomes as more positive when they anticipated high effort (Figure 3). This result suggests that participants experienced relief when high effort was anticipated but avoided.

**Fig 3.**
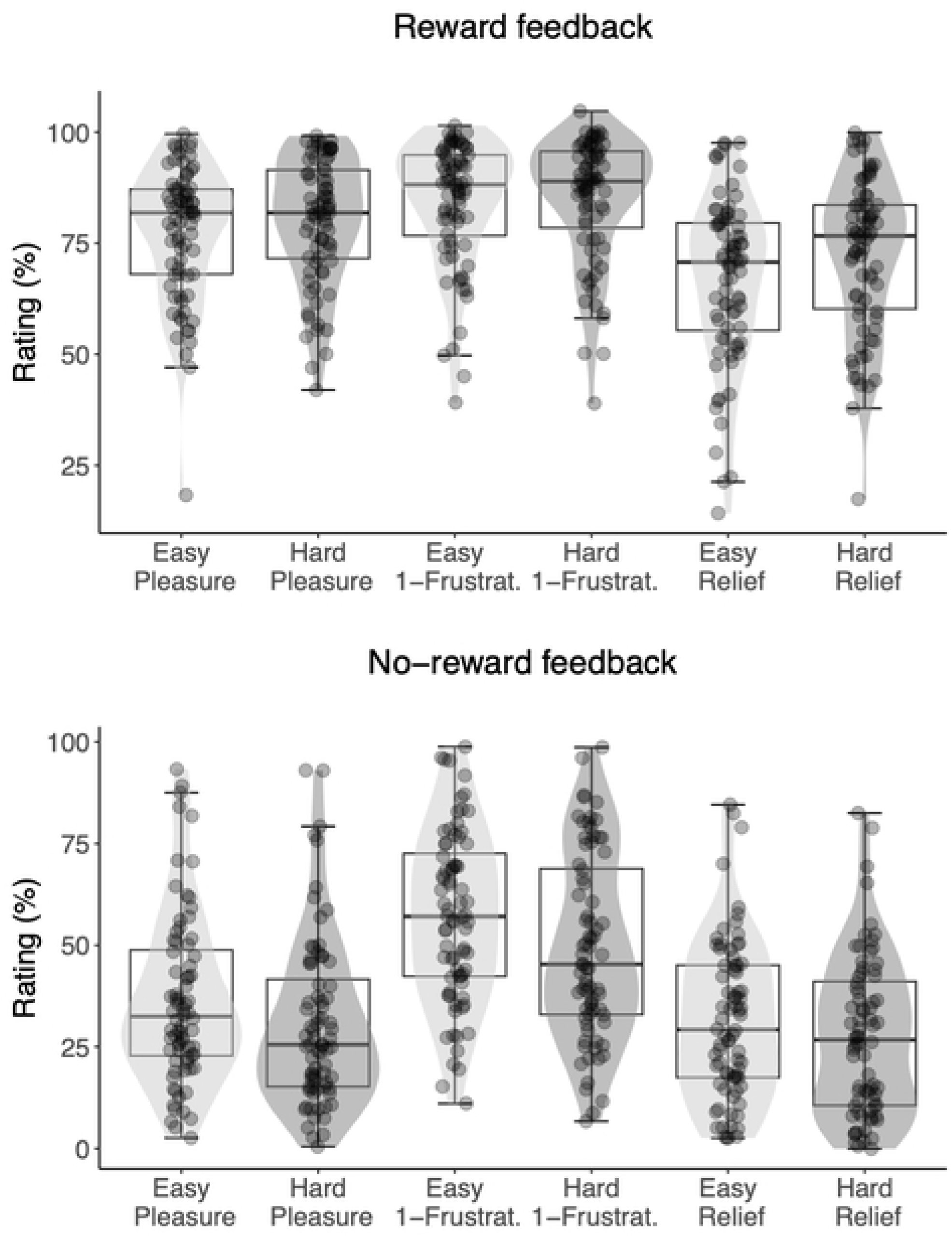
Experiment 2. Ratings of reward-related feedback that followed the gambling task, by difficulty level (easy or hard), and affective dimension (pleasure, reverse-scored frustration, and relief). Each dot represents the subject-level average across 10 repetitions of the rating. For either a reward (top) or no-reward (bottom) outcome, light grey density shades correspond to low effort anticipation and dark grey density shades correspond to high effort anticipation. The horizontal line represents the median, the box represents the interquartile range, and the whiskers extend to the last data point within 1.5 times the interquartile range.

To test the hypothesis that a predisposition towards cognitive effort (as measured with the NFC questionnaire) would reduce this relief effect, we included participants’ NFC score as a continuous predictor in the best-fitting model described above (the two-way model including the interaction between outcome and difficulty level). Under this new three-way interaction model, observed data were more likely compared to the former best model (BF = 2,90 x 10^9^) and the null model (BF = 2.63 x 10^969^). In other words, there was strong evidence that the disposition towards cognitive effort moderated the interaction effect between outcome and difficulty level.

To further explore the effect of NFC on reward processing, we conducted a full Bayesian estimation of parameter values. We fit model 5) with the “brms” R package (46), and we inspected the posterior probability distributions for every interaction parameter that included NFC. Confirming the previous model comparison, we observed a negative three-way interaction *outcome* x *difficulty level* x *NFC* (estimate = -0.15; 95% credible interval = [-0.27 0.04]; posterior probability = 0.98), indicating that the *outcome* x *difficulty level* interaction was moderated by *NFC*. As can be seen in Figure 4B, the differences in subjective ratings between difficulty conditions were attenuated, in both reward conditions, for participants with higher NFC scores. This effect can also be observed in the raw data in Figure 5, where the *outcome* x *difficulty level* crossover interaction is shown for two subsamples of participants scoring at the extremes of the NFC scale. Both the more positive ratings of reward feedback in the difficult condition, as well as the more positive ratings of no reward feedback in the easy condition were attenuated for people in high in NFC.

**Fig 4.**
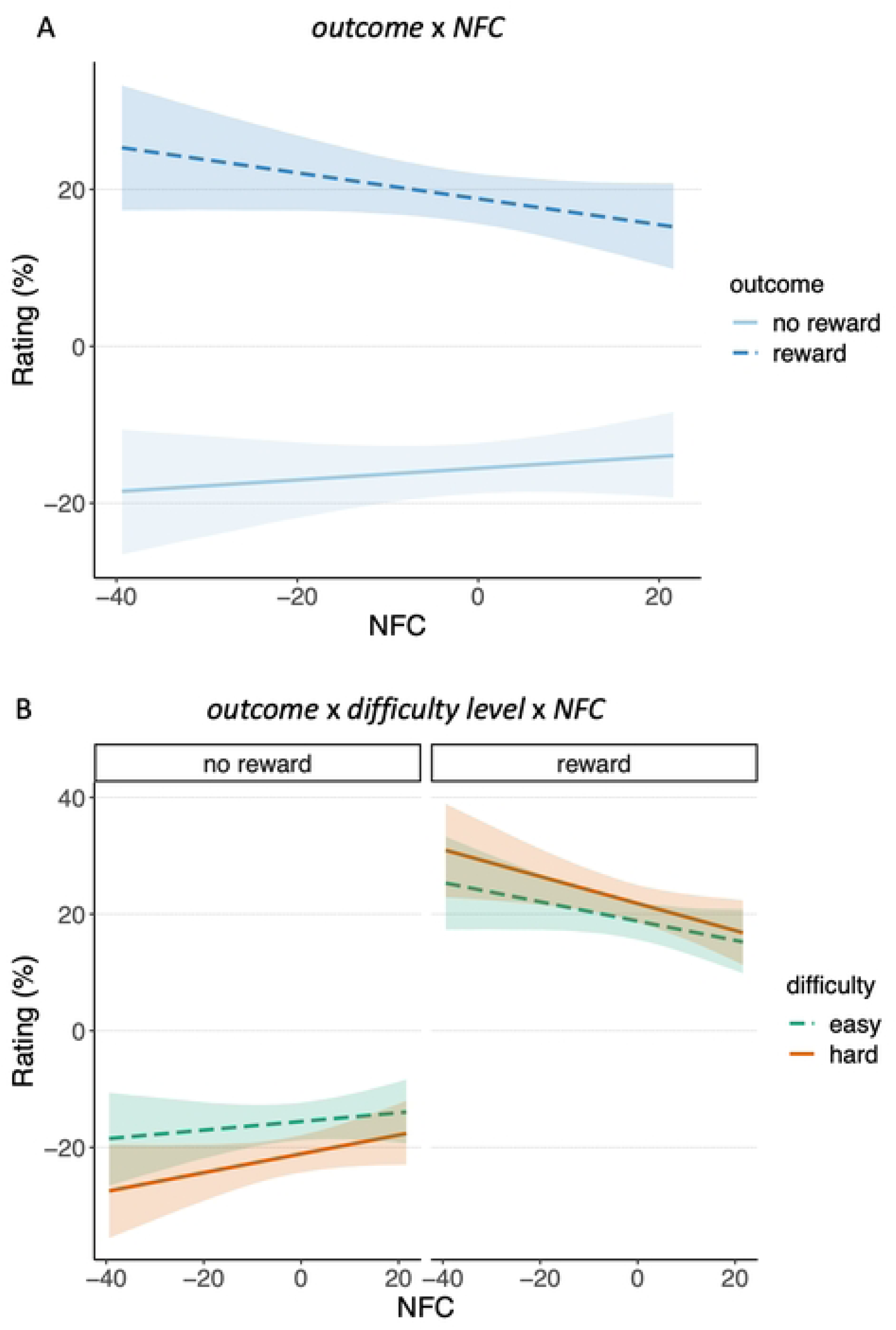
Results of the Bayesian parameter estimation for model 5. Interaction effects of predictors outcome and NFC (panel A) and outcome, difficulty level, and NFC (panel B) on the estimated feedback rating. The regression lines represent the mean of posterior probability samples, for each condition and NFC score. The shading represents the 95% credible interval around them.

**Fig 5.**
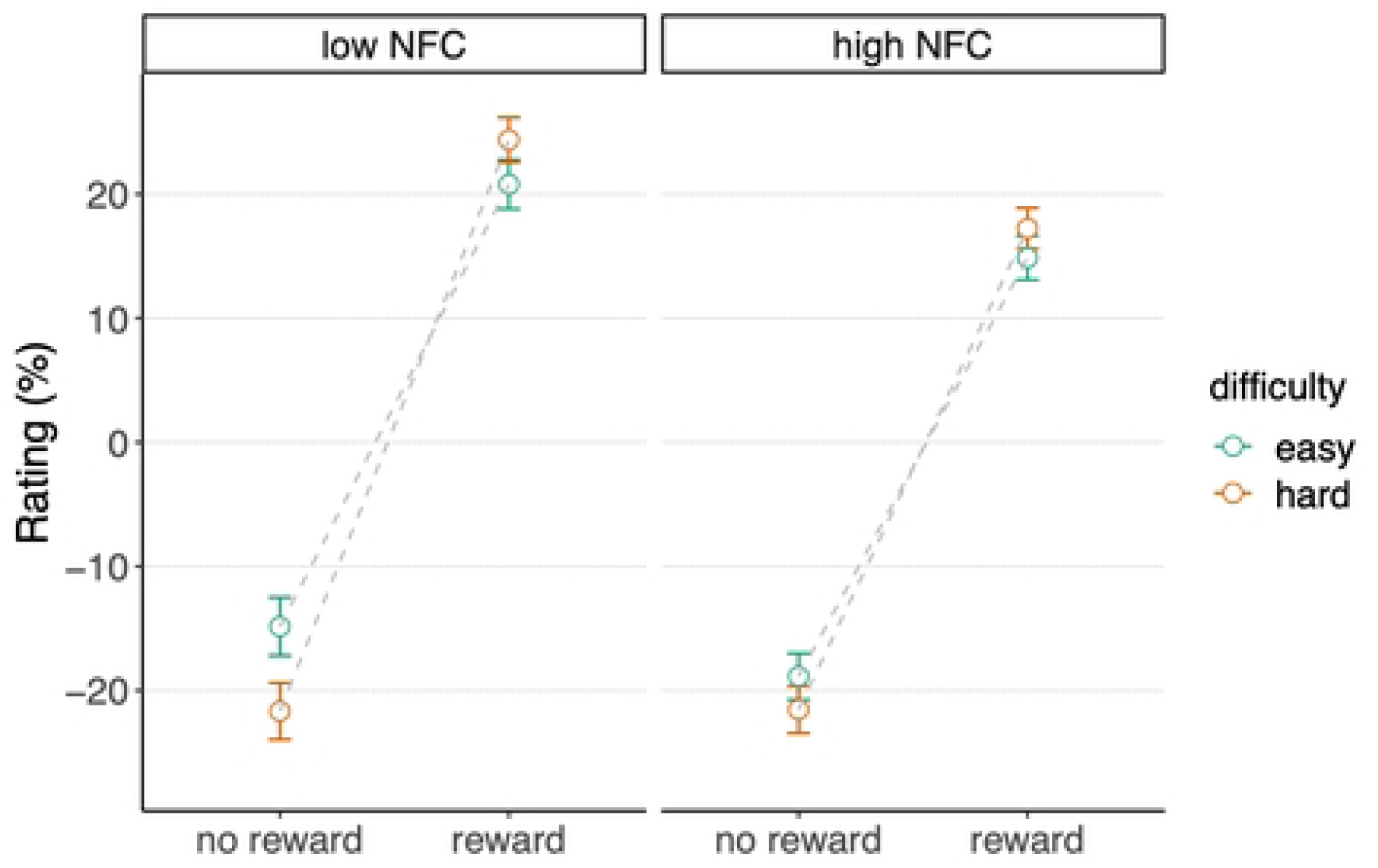
Ratings of reward-related feedback that followed the gambling task. Crossover interaction between outcome and difficulty level for two subsamples of participants with extreme NFC scores. Low NFC = participants within 0-30 percentiles. High NFC = participants within 70-100 percentiles. The three affective dimensions (pleasure, reverse-scored frustration, relief) and the ten repetitions were aggregated. Error bars represent the within-subject confidence intervals. The inter-subject variance has been removed by subtracting the subject mean from each rating, and adding the group mean (68).

Additionally, we observed a negative two-way interaction *outcome* x *NFC* (estimate = -0.24; 95% credible interval = [-0.32 -0.16]; posterior probability = 1), suggesting that the main effect of outcome was also moderated by *NFC*. As can be seen in Figure 4A, participants low on NFC reported more extreme ratings for both reward and no-reward feedback.

To investigate this pattern of results further, we also verified whether NFC predicted cognitive performance in the arithmetic task. We analyzed reaction times and accuracy of the arithmetic task’s responses and compared models including the effects of 1) *difficulty level*, 2) *NFC*, 3) *difficulty level + NFC*, 4) *difficulty level x NFC.* The null model was a simple intercept model, and all models included the random factors *participants* as nuisance. Reaction times (for correct responses) were best explained by the *difficulty level X NFC* interaction model, under which the observed data were BF10 = 1.64 x 10^336^ times more likely to be produced than under the *null* model. This model also explained the observed data BF10 = 8.32 times better than the second-best model, which only included the main effect of *difficulty level*. Similarly, accuracy was best explained by the *difficulty level X NFC* interaction model, under which the observed data were BF10 = 3.50 x 10^74^ times more likely to be produced than under the *null* model. This model explained the observed data BF10 = 8.73 times better than the second-best model, which only included the main effect of *difficulty level*. In other words, for both reaction times and accuracy, there was moderate evidence in support of the hypothesis that NFC interacted with the difficulty level in predicting performance. As can be seen in Figure 6, while the two subsamples of participants scoring at the extremes of the NFC scale showed similar performance for the easy condition, participants scoring high on NFC were more accurate and faster in the hard condition.

**Fig 6.**
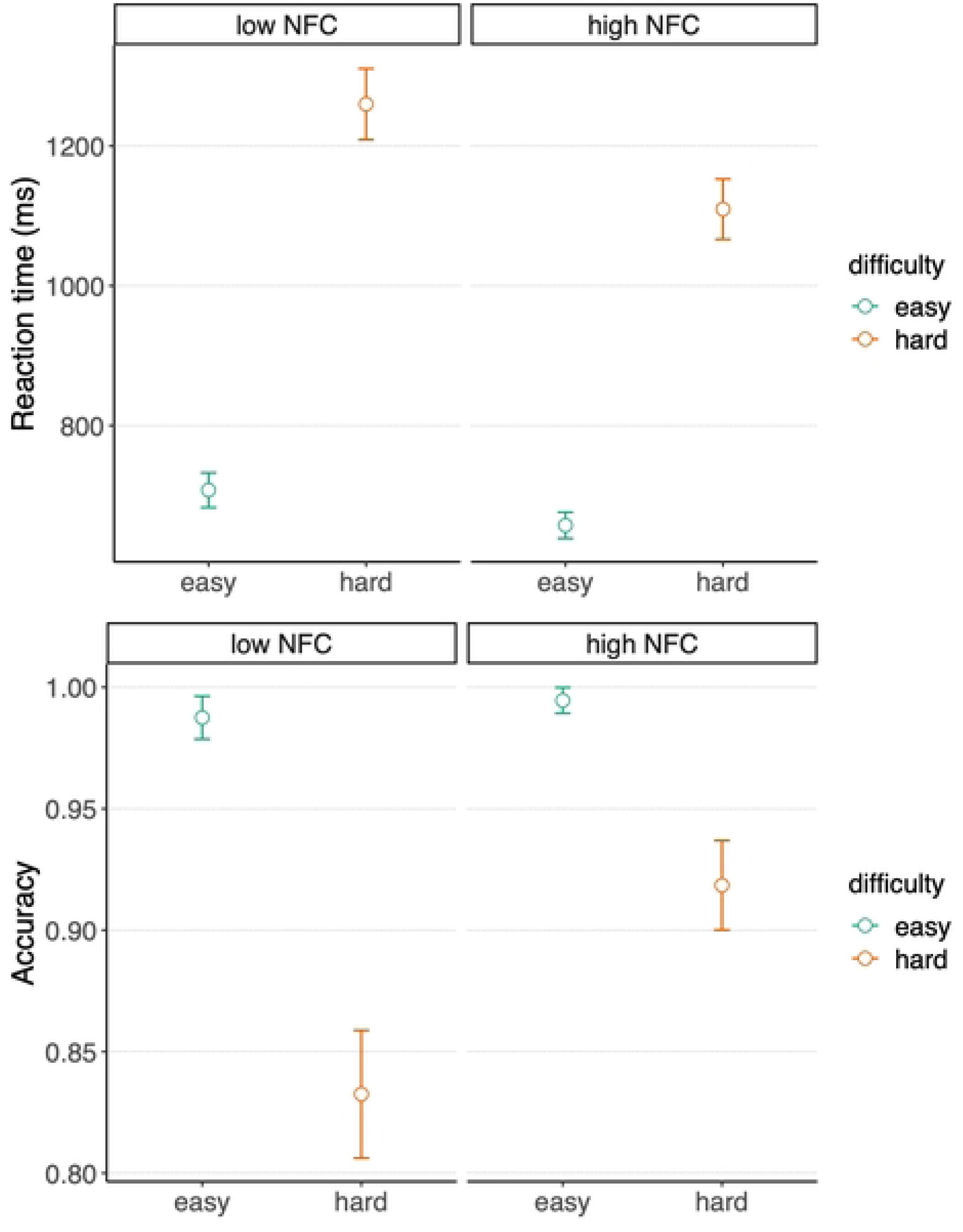
Effect of NFC on arithmetic performance. Reaction times (top) and accuracy (bottom) are reported for two subgroups of participants scoring below the 30^th^ percentile (low NFC, N = 20, NFC < 33) or above the 70^th^ percentile (high NFC, N = 23, NFC > 46.5) of the group distribution of NFC scores, separately for the easy and the hard arithmetic conditions. Error bars represent the within-subject confidence intervals. The inter-subject variance has been removed by subtracting the subject mean from each rating, and adding the group mean (68).

Finally, we analyzed the subjective ratings of the performance feedback (Figure 7). Participants rated positive feedback following the high effort task as more positive compared to following the low effort task. The Bayesian one-tailed t-tests showed strong evidence for this hypothesis (M hard = 74.8, SD = 23.2; M easy = 71.4, SD = 24.3; BF+0 = 6.99 x 10^4^).

**Fig 7.**
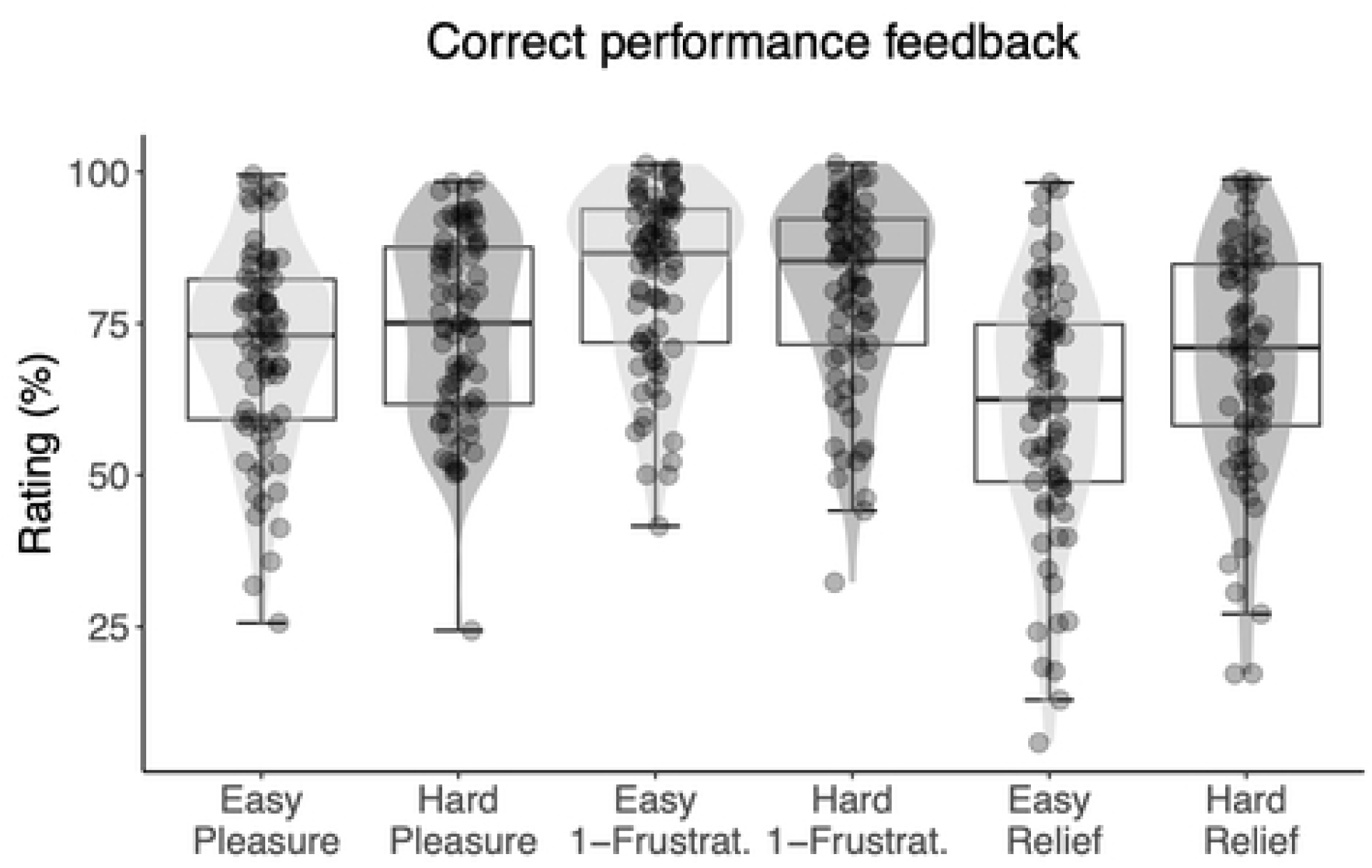
Ratings of performance feedback that followed the effort task, by difficulty level (easy or hard), and affective dimension (pleasure, reverse-scored frustration, and relief). Each dot represents the subject-level average across ∼13 repetitions of the rating. Light grey density shades correspond to correct feedback following low effort and dark grey density shades corresponds to correct feedback following high effort. The horizontal line represents the median, the box represents the interquartile range, and the whiskers extend to the last data point within 1.5 times the interquartile range.

## Discussion

Exerting cognitive effort carries a cost (47). Over the last decade there has been an abundance of empirical evidence that supports this notion, showing that effort exertion is avoided (5,7,8) or traded for reward . Here, by extension, we hypothesized that the avoidance of anticipated cognitive effort carries positive value, because it provides relief from an anticipated cost, which in turn bolsters reward (1,48–54). We tested this hypothesis by adapting a paradigm that we previously used to show that rewards were perceived as more positive when they signaled avoidance of a demanding task (1). However, in this previous experiment, avoidance of the demanding task meant that no task would be performed at all. Therefore, this result could also be driven by opportunity costs (28) rather than effort per se. Here, we adapted this experimental paradigm to overcome this limitation. In our new task, we measured subjective ratings of outcomes that signaled whether an upcoming arithmetic task (29) could be avoided. On each trial, this arithmetic task carried either low or high effort demands, which allowed us to specifically measure the effect of effort context on ratings. In addition, we ran a replication of this study which also investigated whether NFC (32) would act as a moderator on any relief effects. A number of important new results emerged from this study.

First, in both Experiments 1 and 2, the performance measures and subjective ratings suggested that people found the effortful version of the arithmetic task to be more demanding (replicating Vassena et al. (29)). Participants were slower and made more errors on the hard task, and rated it as more difficult and less pleasant. However, at the same time, the motivation to perform both task’s versions was high and similar, and participants also reported the same amount of pleasure when successfully completing it. These results suggest that the hard task yielded more negative ratings because of the costs associated with increased effort exertion (35), yet this negative evaluation did not bias the participants’ motivation. Therefore, they also suggest that people allocated more effort in the hard condition, but that this was not driven by differences in motivation for approaching these tasks, but rather by differences in difficulty.

Second and central to our main hypothesis, we found that rewards were perceived as more positive when they meant people would avoid the harder, more effort-demanding, task. This results extends our prior findings (1), but now more clearly indicating that the avoidance of cognitive effort is perceived as rewarding, translating to a form of relief. Indeed, previous neuroscientific research suggests that the relief of pain involves reward processing pathways (17,18). Our finding also suggests that reward is evaluated in relation to prospective effort. Importantly, because we ensured that opportunity costs (28) were equal between effort conditions, we can be sure that this relief effect is truly driven by effort anticipation.

In both experiments, we also found that effort anticipation influenced ratings of unrewarded outcomes, which signaled that effort could not be avoided. Specifically, people viewed no reward outcomes as more positive if they led to a low-compared to a high-effort arithmetic trial. This result suggests that the prospect of exerting low amounts of effort is perceived as more positive compared to exerting high amounts of effort. This finding is compatible with earlier empirical studies and theoretical models that have associated effort with avoidance and/or reward devaluation/discount (e.g., 2,5,54,55).

Interestingly, in Experiment 2 we found that this relief effect was modulated by participants’ NFC (32). As expected, people with high NFC showed a smaller relief effect. This finding is important because it shows that the cost-benefit analyses carried out during effort allocation do not just consider structural factors (such as reward probability or objective task difficulty), but also specific dispositions or motivational states (55). Therefore, our results inform theoretical models of metacontrol, and of the role of motivation in decision making (57). They clearly signal the need for such models to consider intrinsic disposition towards effortful tasks, and show how they may influence the processing of reward (see 57).

It is tempting to think that the attenuated relief effect for individuals high in NFC suggests that such individuals are not averse to cognitive effort, or perhaps even value its exertion (55). This would pose a challenge to the, presumably universal, law of least mental effort (5). In fact, recent research suggests that explicitly rewarding the selection of high-effort actions biases people to choose more effortful lines of action (59,60). However, an increased NFC may simply reflect a heightened subjective value of succeeding at demanding tasks, or of the rewards obtained after demanding tasks. Alternatively, people with high NFC may believe that completing hard tasks demonstrates self-efficacy or competence. Under this view, these factors would offset the intrinsic cost of effort during decision making (61). In other words, a NFC may reflect heightened stakes for cognitive success, rather than lower cognitive costs. Future research may specifically address these open questions by probing in a more granular fashion which of these factors affect effort-based decision making. Finally, we hasten to mention that people high on NFC showed an attenuated relief effect, but not a reversed one. This finding is not inconsistent with the idea that these people simply carry a smaller, but still positive, cost of mental effort.

Interestingly, the results from Experiment 2 also revealed that participants high on NFC solved the hard arithmetic task faster and better than those low on this dimension. This result appears to be at odds with previous studies available in the literature. A range of studies have not found any systematic relationship between the NFC and behavioral performance (using a variety of tasks that demand cognitive control functions such as conflict processing or response inhibition; (62)). Accordingly, some caution is needed when interpreting the modulatory influence of the NFC on the relief effect. While it is possible that this effect is driven by an increased intrinsic motivation to engage in effortful tasks (as explained above), it is also possible that it was caused by the level of required effort being “objectively” lower for participants high on NFC. In other words, the increased task performance on the hard arithmetic task by this group of participants may reflect that it was not difficult enough for them, leading to an attenuated relief effect.

Of course, the current study does not allow us to adjudicate between these two competing, but not mutually exclusive, accounts. That being said, we would like to note that the questionnaire was administered after task completion, so it is not possible that it induced biases or expectations about effort exertion that influenced behavior on the task. More generally, the correlational nature of this finding prevents us from drawing any causal inferences between the relief effect and NFC. Hence, additional studies are needed to determine the mechanistic nature of the modulation of the relief effect by NFC. For example, one group of participants may be trained to value effort using one of the manipulations introduced by Clay et al. (58) and Lin et al. (59). If our results were truly driven by altered representations of effort costs, then this group should also show a similar attenuation of the relief effect.

Finally, in Experiment 2 we also found that participants valued positive feedback after the arithmetic task more highly when they had successfully performed the hard compared to easy task. This result suggests that effort expenditure (as opposed to anticipation only) can also influence reward processing (see also 62,63), retrospectively increasing the value of rewards when more effort is exerted. This finding bears resemblance to the IKEA effect (55,65), according to which people value a product more if it is produced by their own efforts compared to the same product acquired by other means. In addition, it calls to mind the idea that self-efficacy (66), typically achieved through effort, carries value, and contributes to a positive sense of competence (c.f. self-determination theory (67)).

Some methodological limitations warrant a final note. The current studies used a relatively small sample of student participants. This is especially noteworthy for Experiment 2, where we aimed to assess associations between individual differences. Our results invite a replication with a larger and more heterogeneous sample. Second, our effects emerged in the context of a simple gambling task, and so it is unknown whether the relief effect would emerge in other contexts. For example, what would happen if reward feedback, unlike the current studies, would be useful for decision making (e.g., in the context of reinforcement learning)? Third, we established a robust relief effect based only on subjective ratings of the reward-related feedback, but it is unclear whether this novel paradigm would still elicit its neural correlates (1). Neuroimaging work using electrophysiology would not only be useful for validating the relief effect at the single trial level, but also for developing formal neurocomputational models of cost-benefit tradeoffs in effort allocation.

In sum, this study provides novel evidence for the notion that cognitive effort carries a cost, and that this cost is tightly linked to both reward processing and cost-benefit analyses of effort allocation. More precisely, our results suggest that if anticipated effort is avoided, it is returned as a reward, and experienced as relief. Intriguingly, this effect is attenuated in people who report being more disposed towards effortful tasks, suggesting they are less sensitive to cognitive costs. This raises the outstanding question of whether differences in NFC mitigate the aversive nature of cognitive effort, or whether they result in heightened stakes of engaging with, and succeeding at, hard cognitive tasks.

## References

1. Gheza D, De Raedt R, Baeken C, Pourtois G. Integration of reward with cost anticipation during performance monitoring revealed by ERPs and EEG spectral perturbations. NeuroImage. 2018;173:153–64.

2. Shenhav A, Botvinick M, Cohen J. The expected value of control: An integrative theory of anterior cingulate cortex function. Neuron. 2013;79(2):217–40.

3. Silvestrini N, Musslick S, Berry AS, Vassena E. An integrative effort: Bridging motivational intensity theory and recent neurocomputational and neuronal models of effort and control allocation. Psychol Rev. 2022;

4. Hull C. Principles of Behavior. New York, USA: Appleton-Century-Crofts; 1943.

5. Kool W, McGuire JT, Rosen ZB, Botvinick MM. Decision Making and the Avoidance of Cogntive Demand. J Exp Psychol Gen. 2010;139(4):665–82.

6. Bustos B, Colvett JS, Bugg J, Kool W. Humans do not avoid reactively implementing cognitive control [Internet]. 2023 [cited 2023 Jun 13]. Available from: https://doi.org/10.31234/osf.io/x5vgj

7. Schouppe N, Ridderinkhof KR, Verguts T, Notebaert W. Context-specific control and context selection in conflict tasks. Acta Psychol (Amst). 2014 Feb 1;146:63–6.

8. Sayalı C, Badre D. Neural systems of cognitive demand avoidance. Neuropsychologia. 2019;123(October 2017):41–54.

9. Sayali C, Rubin-McGregor J, Badre D. Policy abstraction as a predictor of cognitive effort [Internet]. PsyArXiv; 2022 Jun [cited 2023 Jun 13]. Available from: https://osf.io/by7gk

10. Vogel TA, Savelson ZM, Otto AR, Roy M. Forced choices reveal a trade-off between cognitive effort and physical pain. eLife. 2020 Nov 17;9:e59410.

11. Cohen JD. Cognitive Control. In: The Wiley Handbook of Cognitive Control [Internet]. John Wiley & Sons, Ltd; 2017 [cited 2023 Jun 12]. p. 1–28. Available from: https://onlinelibrary.wiley.com/doi/abs/10.1002/9781118920497.ch1

12. Miller EK, Cohen JD. An integrative theory of prefrontal cortex function. 2001;167–202.

13. Shenhav A, Musslick S, Lieder F, Kool W, Griffiths TL, Cohen JD, et al. Toward a Rational and Mechanistic Account of Mental Effort. Annu Rev Neurosci. 2017;40(1):annurev-neuro-072116-031526.

14. Botvinick MM, Huffstetler S, McGuire JT. Effort discounting in human nucleus accumbens. Cogn Affect Behav Neurosci. 2009;

15. Massar SAA, Libedinsky C, Weiyan C, Huettel SA, Chee MWL. Separate and overlapping brain areas encode subjective value during delay and effort discounting. NeuroImage. 2015 Oct 15;120:104–13.

16. Salamone JD, Correa M. The Mysterious Motivational Functions of Mesolimbic Dopamine. Neuron. 2012;76(3):470–85.

17. Leknes S, Lee M, Berna C, Andersson J, Tracey I. Relief as a Reward: Hedonic and Neural Responses to Safety from Pain. PLOS ONE. 2011 Apr 7;6(4):e17870.

18. Porreca F, Navratilova E. Reward, motivation, and emotion of pain and its relief. Pain. 2017 Apr;158 Suppl 1(Suppl 1):S43–9.

19. Hajcak G, Holroyd CB, Moser JS, Simons RF. Brain potentials associated with expected and unexpected good and bad outcomes. Psychophysiology. 2005;42(2):161–70.

20. Klein ED, Bhatt RS, Zentall TR. Contrast and the justification of effort. Psychon Bull Rev. 2005;12(2):335–9.

21. Proudfit GH. The reward positivity: From basic research on reward to a biomarker for depression. Psychophysiology. 2015;52(4):449–59.

22. Webb CA, Auerbach RP, Bondy E, Stanton CH, Foti D, Pizzagalli DA. Abnormal neural responses to feedback in depressed adolescents. J Abnorm Psychol. 2017;126(1):19–31.

23. Cohen MX, Elger CE, Ranganath C. Reward expectation modulates feedback-related negativity and EEG spectra. NeuroImage. 2007;35(2):968–78.

24. Marco-Pallares J, Cucurell D, Cunillera T, Garc??a R, Andr??s-Pueyo A, M??nte TF, et al. Human oscillatory activity associated to reward processing in a gambling task. Neuropsychologia. 2008;46(1):241–8.

25. Mas-herrero E, Ripollés P, Hajihosseini A, Rodríguez-fornells A, Marco-pallarés J. NeuroImage Beta oscillations and reward processing : Coupling oscillatory activity and hemodynamic responses. NeuroImage. 2015;119:13–9.

26. Botvinick MM, Braver TS. Motivation and Cognitive Control: From Behavior to Neural Mechanism. Annu Rev Psychol. 2015;66(1):83–113.

27. Westbrook A, Braver TS. Dopamine Does Double Duty in Motivating Cognitive Effort. Neuron. 2016;89(4):695–710.

28. Kurzban R, Duckworth A, Kable JW, Myers J. An opportunity cost model of subjective effort and task performance. Behav Brain Sci. 2013;36(06):661–79.

29. Vassena E, Silvetti M, Boehler CN, Achten E, Fias W, Verguts T. Overlapping neural systems represent cognitive effort and reward anticipation. PLoS ONE. 2014;9(3).

30. Bogdanov M, Nitschke JP, LoParco S, Bartz JA, Otto AR. Acute Psychosocial Stress Increases Cognitive-Effort Avoidance. Psychol Sci. 2021 Sep 1;32(9):1463–75.

31. Pizzagalli DA. Depression, stress, and anhedonia: toward a synthesis and integrated model. Annu Rev Clin Psychol. 2014;10(January):393–423.

32. Cacioppo JT, Petty RE. The need for cognition. J Pers Soc Psychol. 1982;42:116–31.

33. Zerna J, Strobel A, Strobel A. The role of Need for Cognition in well-being – Review and meta-analyses of associations and potentially underlying mechanisms [Internet]. PsyArXiv; 2021 [cited 2023 Jun 4]. Available from: https://psyarxiv.com/p6gwh/

34. Cacioppo JT, Petty RE, Feinstein JA, Jarvis WBG. Dispositional differences in cognitive motivation: The life and times of individuals varying in need for cognition. Psychol Bull. 1996;119(2):197–253.

35. Yee DM, Leng X, Shenhav A, Braver TS. Aversive motivation and cognitive control. Neurosci Biobehav Rev. 2022 Feb;133:104493.

36. Hajcak G, Moser JS, Holroyd CB, Simons RF. It’s worse than you thought: The feedback negativity and violations of reward prediction in gambling tasks. Psychophysiology. 2007;44(6):905–12.

37. Vassena E, Cobbaert S, Andres M, Fias W, Verguts T. Unsigned value prediction-error modulates the motor system in absence of choice. NeuroImage. 2015;122:73–9.

38. Vassena E, Deraeve J, Alexander WH. Task-specific prioritization of reward and effort information: Novel insights from behavior and computational modeling. Cogn Affect Behav Neurosci. 2019;

39. Vassena E, Gerrits R, Demanet J, Verguts T, Siugzdaite R. Anticipation of a mentally effortful task recruits Dorsolateral Prefrontal Cortex: An fNIRS validation study. Neuropsychologia. 2018;123(October 2017):106–15.

40. Cacioppo, J. T., Petty, R. E., Kao, C. F. Need for Cognition Scale. Measurement Instrument Database for the Social Science. [Internet]. 2013 [cited 2023 May 23]. Available from: Retrieved from www.midss.org

41. Psychology Software Tools, Inc. [E-Prime 2.0] [Internet]. Sharpsburg, PA: Psychology Software Tools, Inc.; 2008. Available from: https://support.pstnet.com/.

42. Rouder JN, Morey RD, Verhagen J, Swagman AR, Wagenmakers EJ. Bayesian analysis of factorial designs. Psychol Methods. 2017;22(2):304–21.

43. Schindler S, Schettino A, Pourtois G. Electrophysiological correlates of the interplay between low-level visual features and emotional content during word reading. Sci Rep. 2018;

44. R Core Team. R. R Core Team. 2017.

45. Morey RD, Rouder JN. Bayesfactor: Computation of Bayes factors for common designs. R package version 0.9.12-2. BayesFactor Comput Bayes Factors Common Des. 2015;

46. Bürkner PC. brms : An R Package for Bayesian Multilevel Models Using Stan. J Stat Softw. 2017;80(1):1–28.

47. Kool W, Botvinick M. Mental labour. Nat Hum Behav. 2018;2(12):899–908.

48. Devine S, Vassena E, Otto AR. More than a feeling: physiological measures of affect index the integration of effort costs and rewards during anticipatory effort evaluation. Cogn Affect Behav Neurosci [Internet]. 2023 Apr 14 [cited 2023 Jun 13]; Available from: https://link.springer.com/10.3758/s13415-023-01095-3

49. Kool W, Gershman SJ, Cushman FA. Cost-Benefit Arbitration Between Multiple Reinforcement-Learning Systems. Psychol Sci. 2017;28(9):1321–33.

50. Kool W, Shenhav A. Cognitive Control as Cost-Benefit Decision Making. 2017;

51. Padmala S, Pessoa L. Reward Reduces Conflict by Enhancing Attentional Control and Biasing Visual Cortical Processing. J Cogn Neurosci. 2011 Nov 1;23(11):3419–32.

52. Treadway MT, Buckholtz JW, Schwartzman AN, Lambert WE, Zald DH. Worth the “EEfRT”? The effort expenditure for rewards task as an objective measure of motivation and anhedonia. PLoS ONE. 2009;4(8):1–9.

53. Westbrook A, Kester D, Braver TS. What Is the Subjective Cost of Cognitive Effort? Load, Trait, and Aging Effects Revealed by Economic Preference. PLoS ONE. 2013;8(7):1–8.

54. Westbrook A, Van Den Bosch R, Määttä JI, Hofmans L, Papadopetraki D, Cools R, et al. Dopamine promotes cognitive effort by biasing the benefits versus costs of cognitive work. Science. 2020 Mar 20;367(6484):1362–6.

55. Inzlicht M, Shenhav A, Olivola CY. The Effort Paradox: Effort Is Both Costly and Valued. Trends Cogn Sci. 2018;22(4):337–49.

56. McGuire JT, Botvinick MM. Prefrontal cortex, cognitive control, and the registration of decision costs. Proc Natl Acad Sci. 2010;107(17):7922–6.

57. Vassena E, Deraeve J, Alexander WH. Predicting Motivation: Computational Models of PFC Can Explain Neural Coding of Motivation and Effort-based Decision-making in Health and Disease. J Cogn Neurosci. 2017;29(10):1633–45.

58. Thompson EP, Chaiken S, Hazlewood JD. Need for cognition and desire for control as moderators of extrinsic reward effects: a person x situation approach to the study of intrinsic motivation. J Pers Soc Psychol. 1993 Jun;64(6):987–99.

59. Clay G, Mlynski C, Korb FM, Goschke T, Job V. Rewarding cognitive effort increases the intrinsic value of mental labor. Proc Natl Acad Sci. 2022 Feb;119(5):e2111785119.

60. Lin H, Westbrook A, Fan F, Inzlicht M. An experimental manipulation increases the value of effort [Internet]. PsyArXiv; 2021 Feb [cited 2023 Jun 13]. Available from: https://osf.io/gnk4m

61. Kool W, Botvinick M. The intrinsic cost of cognitive control. Behav Brain Sci. 2013 Dec;36(6):697– 8.

62. Gärtner A, Grass J, Wolff M, Goschke T, Strobel A, Strobel A. No relation of Need for Cognition to basic executive functions. J Pers. 2021 Dec;89(6):1113–25.

63. Bogdanov M, Renault H, LoParco S, Weinberg A, Otto AR. Cognitive effort exertion enhances electrophysiological responses to rewarding outcomes. Cereb Cortex. 2022 Sep 19;32(19):4255– 70.

64. Ma Q, Meng L, Wang L, Shen Q. I endeavor to make it: Effort increases valuation of subsequent monetary reward. Behav Brain Res. 2013;261:1–7.

65. Norton MI, Mochon D, Ariely D. The IKEA effect: When labor leads to love. J Consum Psychol. 2012 Jul 1;22(3):453–60.

66. Bandura A. Self-efficacy: toward a unifying theory of behavioral change. Psychol Rev. 1977 Mar;84(2):191–215.

67. Ryan RM, Deci EL. Intrinsic and Extrinsic Motivations : Classic Definitions and New Directions. 2000;67:54–67.

68. Morey RD. Confidence Intervals from Normalized Data: A correction to Cousineau (2005). Tutor Quant Methods Psychol. 2008 Sep 1;4(2):61–4.

